# Resource-dependent heterosynaptic spike-timing-dependent plasticity in recurrent networks with and without synaptic degeneration

**DOI:** 10.1101/2025.03.16.643509

**Authors:** James Humble

## Abstract

Many computational models that incorporate spike-timing-dependent plasticity (STDP) have shown the ability to learn from stimuli, supporting theories that STDP is a sufficient basis for learning and memory. However, to prevent runaway activity and potentiation, particularly within recurrent networks, additional global mechanisms are commonly necessary. A STDP-based learning rule, which involves local resource-dependent potentiation and heterosynaptic depression, is shown to enable stable learning in recurrent spiking networks. A balance between potentiation and depression facilitates synaptic homeostasis, and learned synaptic characteristics align with experimental observations. Furthermore, this resource-based STDP learning rule demonstrates an innate compensatory mechanism for synaptic degeneration.

## 1 Introduction

The concept of spike-timing-dependent plasticity (STDP) has been thoroughly researched and frequently serves as a foundation for learning in computational models. Various studies adopt STDP in diverse formats. For instance, it may be utilized with either an additive or a multiplicative rule for updates: Potentiation or depression may depend on or be independent of a synapse’s weight. Different STDP implementations can lead to varied outcomes, with some rules more closely reflecting phenomena observed in experiments. These variations include 1) synaptic weight distributions, 2) the presence of nonpotentiable synapses, 3) silent synapses, 4) synaptic persistence, and 5) competition between synapses:

1. Empirically identified synaptic weight distributions generally display a uni-modal pattern that peaks close to zero, characterized by numerous weak synapses and a few strong connections forming a long tail (Buzski, 2004; Yasumatsu et al., 2008; Kasai, 2023). In computational models, synaptic distributions change based on whether STDP is implemented additively or multiplicatively. In feedforward networks, additive STDP often produces bi-modal distributions, with peaks located near zero and the upper limit of synaptic weight (Rossum et al., 2000; Barbour et al., 2007; Morrison et al., 2008). In contrast, multiplicative STDP often generates uni-modal distributions with a peak situated between zero and the upper bound (Rossum et al., 2000; Barbour et al., 2007; Morrison et al., 2008). In recurrent networks, additive STDP can produce a uni-modal distribution with a peak at the upper-bound and multiplicative, uni- or multi-modal distributions (Morrison et al., 2007).
2. Silent synapses are primarily defined by their absence of *α*-amino-3-hydroxy-5-methyl-4-isoxazolepropionic acid (AMPA) receptors, as comprehensively discussed by Montgomery and colleagues (2004). Despite the scarcity of AMPA receptors, these synapses often retain a degree of plasticity due to the presence of *N* -methyl-D-aspartate (NMDA) receptors (Kim et al., 2025). In a study by Brunel et al. (2004), a prominent and sharply delineated subset of silent synapses was integrated into an empirically derived distribution by assessing those potentially undetected because of technological limitations and their deficiency in AMPA receptors. Such a peak is observed in spine volume (Yasumatsu et al., 2008) and synaptic efficacy (Barbour et al., 2007).
3. Research has indicated that certain synapses may not be capable of potentiation. For instance, Debanne et al. (1999) reported the inability to induce potentiation in 24 % of the synapses examined. Debanne and colleagues propose that this nonpotentiability might be attributed to synapses individually reaching saturation. Conversely, computational models often employ either a universal upper limit across all synapses or a global normalizing mechanism and neglecting these constraints may lead to excessive activity and potentiation (Rossum et al., 2000).
4. The persistence of synapses is typically considered to be fundamentally important for memory. In their study, Billings et al. (2009) investigated the durability of synaptic weights governed by STDP principles and found that additive STDP facilitates stability, whereas multiplicative STDP causes instability due to rapid weight variations. Empirical evidence from long-term potentiation (LTP) studies indicates a two-phase persistence: an initial phase that diminishes swiftly and seems to be reliant on neuronal activity (Dong et al., 2015), and a later phase capable of sustaining LTP over prolonged periods, possibly extending to a year (Abraham et al., 2002). There is, however, strong evidence demonstrating that spine volumes fluctuate in the absence of activity and plasticity (reviewed by Kasai (2023)) and that such fluctuations combined with STDP can be stable (Humble et al., 2019). In this case, the mean spine volume of a group of neurons must be persistent rather than the individual synaptic strengths.
5. Whereas STDP following an additive rule is known for its strong competitive interactions, STDP governed by a multiplicative rule typically exhibits limited competition, prompting the incorporation of supplementary mechanisms such as synaptic scaling or intrinsic fluctuations to achieve the competition essential for learning (Rossum et al., 2000; Humble et al., 2019). The inherently competitive aspect of additive STDP typically necessitates enforcing a stringent upper limit on synaptic strength to control excessive potentiation. In contrast, multiplicative STDP operates under more flexible upper limits determined by weight dependency. A drawback of deploying global limits is their assignment of preset values before the learning process, which may not align with biological realism. Moreover, as mentioned above, Debanne et al. (1999) demonstrated the possibility of individual synaptic upper limits.

Furthermore, to control runaway activity, mechanisms such as synaptic scaling (Turrigiano, 2008), inhibition (Bannon et al., 2020; Eckmann et al., 2024), or the Beinenstock-Cooper-Munro rule (Cooper and Bear, 2012) are typically included with Hebbian-based learning rules in recurrent networks. However, these can operate on a slower timescale than STDP.

The above picture is further complicated by experiments showing that heterosynaptic plasticity can occur at unstimulated spines near a stimulated one (reviewed by Chater and Goda (2021)). Essentially, given some homosynaptic activity in a subset of spines, heterosynaptic changes have been observed at unstimulated ones. The distance dependence and direction of these heterosynaptic changes are potentially competitive (Chater et al., 2024).

Motivated by these experimental results, this paper explores ongoing research into computational disparities by utilizing a learning methodology that integrates ‘resources’ alongside heterosynaptic plasticity in a spiking recurrent network. In a computational model of individual neurons (Chen et al., 2013) and a feedforward network (Chapter 5 of (Humble, 2013)), it has previously been shown that heterosynaptic plasticity can be beneficial in controlling activity and plasticity.

In addition to networks with static connectivity, progressive loss of synapses is a hallmark of many neurodegenerative diseases, including Huntington’s, Parkinson’s, and Alzheimer’s (Herms and Dorostkar, 2015; Meftah and Gan, 2023), and neuropsychiatric disorders such as schizophrenia and depression (Penzes et al., 2011). To counteract synaptic loss, the existence of compensatory mechanisms has been suggested that include enlargement of the remaining spines and increased spinogenesis (Bhembre et al., 2023).

By advancing STDP through the incorporation of limited resources for potentiation and merging it with heterosynaptic depression that affects neighboring synapses, the results reveal that this model effectively resolves the discrepancies mentioned above while presenting an innate mechanism for both synaptic homeostasis and synaptic degeneration compensation.

## 2 Methods

The network structure, Fig. 1a, consists of a pool of *N*_exc_ = 200 excitatory neurons recurrently connected with plastic excitatory synapses with a probability of 25 %. Independent Poisson spike trains are presented to the pool with an input connection probability of 10 % from *N*_in_ = 50 spike trains. All connections have axonal delays drawn from a uniform distribution from 1 ms to 5 ms (Lemarchal et al., 2021).

**Figure 1:**
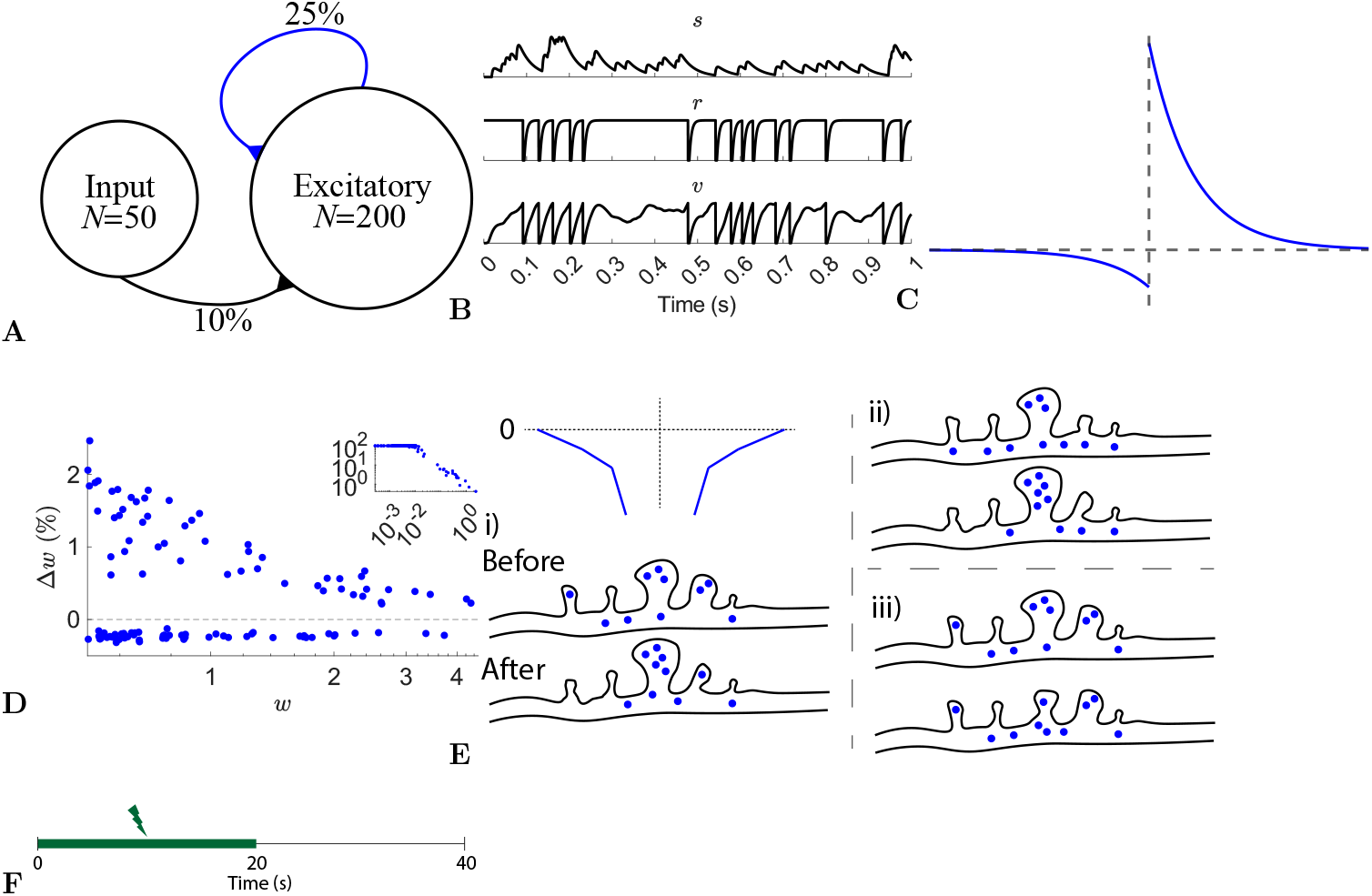
Network, neuron, and synaptic models and stimulation protocol. **(A)** A pool of 200 excitatory neurons receive input from 50 independent Poisson processes. All recurrent synapses undergo STDP. **(B)** A synaptic input, *s*, and one neuron’s refractoriness, *r*, and membrane potential, *v*. **(C)** STDP learning window. **(D)** Synaptic change dependence on weight. **(E)** Three different scenarios for heterosynaptic plasticity: i) potentiation accompanied by heterosynaptic depression of resources (blue circles) from neighbors, ii) potentiation with resources from the pool, and iii) depression. **(F)** Input is provided for the first 20 s.

Excitatory neurons are refractory leaky integrate-and-fire cells, Eqs. 1. The membrane potential of each neuron, *v*, follows low-pass dynamics with a time constant *τ*_*v*_ = 25 ms (Rall, 1969) and is reset when it reaches the firing threshold, *θ* = 1, Fig. 1b bottom.

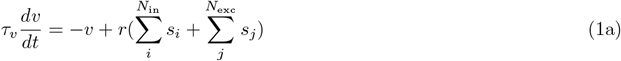

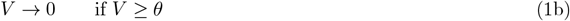

An absolute refractory period of 3 ms is followed by a relative refractoriness period, *τ*_*r*_ = 5 ms, Eq. 2 and Fig. 1b middle.

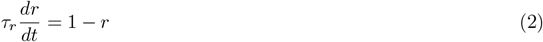

All input and recurrent afferents were modeled as synaptic currents, *s*^*l*^ and *s*, Eqs. 3 and Fig. 1b top. *τ*_*sr*_ = 2.6 ms is the time constant for the synaptic current rise and *τ*_*sf*_ = 31.3 ms the decay constant (Hunt et al., 2022). The input of each synapse consists of binary neuron spikes, *I*_*i*_, scaled by the synapse’s weight, *w*_*i*_.

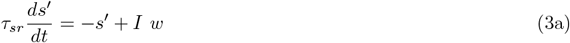

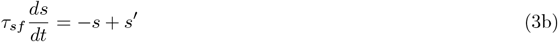

Pairs of presynaptic and postsynaptic spikes evoke changes in synaptic weight, Δ*w* = *f* (*τ*), given as a function of their temporal distance, *τ* = *post-spike time − pre-spike time*, Fig. 1c. Equation 4 describes the STDP function used, where *τ*_*ST DP*_ = 20 ms (Bi and Poo, 1998). Depression is weight dependent, as observed by Bi and Poo (1998), Fig. 1d. Recurrent weights were bounded [0, +*∞*). See below for a description of the inclusion of 0.18 in depression.

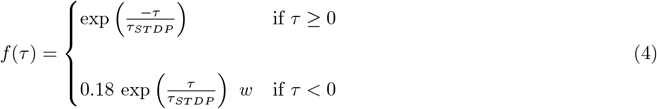

The initial input weights are set with random values drawn from a uniform distribution between 0 and 1. All recurrent weights, *w*, are initially assigned 0 as akin to a newly formed network.

This typical STDP implementation is extended as in Humble (2013) to include the requirement of resources for potentiation paired with potentiation-driven heterosynaptic depression in neighboring synapses. Firstly, a pool of resources is included, *p*. Similarly to many receptors and proteins that undergo degradation or recycling, the pool’s resources decay with a time constant of *τ*_*p*_ = 10 s, Eq. 5.

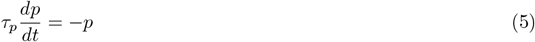

The amounts of the initial resource pool are assigned by Eq. 6 where *ξ* is a random number from a standard normal distribution. The constant multiple was found through a parameter search to ensure that the weights were strong enough that the activity continued after stimulation stopped. A lognormal distribution is chosen as skewed distributions are observed for many aspects of brain dynamics (Buzski and Mizuseki, 2014).

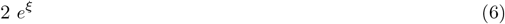

When updating the weights with resource-dependent STDP, there are three scenarios, Fig. 1e:

- When potentiating a synapse, its neighbors are actively depressed by an exponential function of their distance to the potentiating synapse, and the potentiating amount to a maximum distance of 3 synapses.
- When a potentiating synapse and its neighbors have no resources left, the potentiating synapse acquires resources from the pool, if available.
- When a synapse is depressed, its resources are relocated to the pool for reuse.

For simplicity, time constants for resource mobility are neglected; hence, resources can transfer between a synapse and the pool, and vice versa, within a single time step.

With local heterosynaptic plasticity, depression dominates because potentiation events are accompanied by depression, and thus depression has to be decreased. It has previously been found that 0.18 was a suitable multiple for depression such that potentiation and depression were balanced (Humble, 2013). This reduced depression, Fig. 1c, matches the experimental observations of decreased depression relative to potentiation (Bi and Poo, 1998).

The rate of the input spike trains is 50 Hz for the first 20 s, Fig. 1f.

All simulations used forward Euler integration with a time step of *dt* = 0.1 ms and were implemented in MATLAB.

## 3 Results

During stimulation of a typical network, most neurons in the excitatory pool fire with increased firing rates, Figs. 2a and 2b. After stimulation, a subset continues to fire representing a learned memory. Not all neurons are recruited into the memory, due to initial random input and recurrent connectivity and random activity-driven plasticity. Furthermore, after learning, the firing rates of many neurons change, Fig. 2c; nevertheless, the memory is preserved.

**Figure 2:**
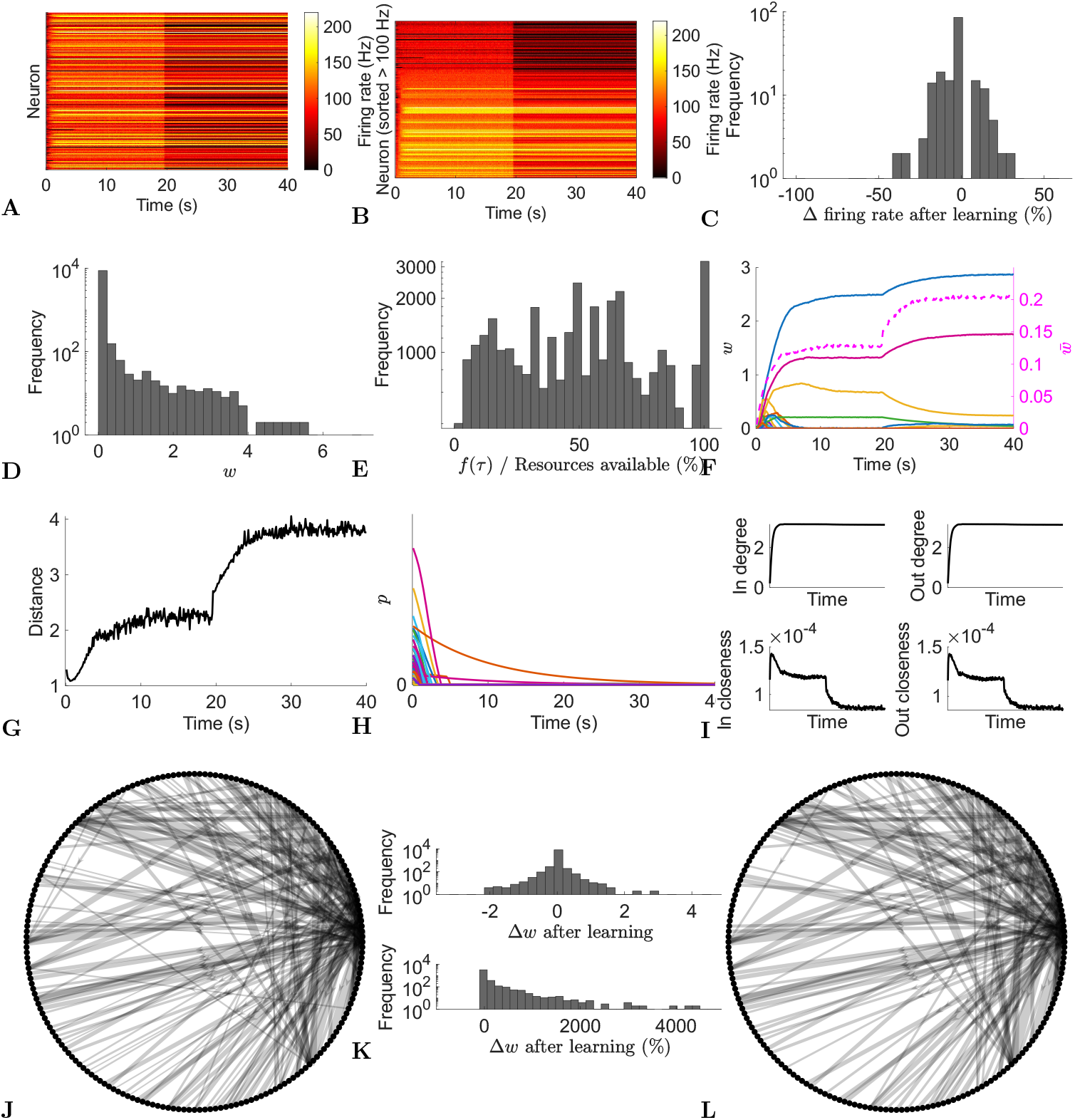
Typical network dynamics. **(A)** Firing rates of the network’s neurons. **(B)** Sorted firing rate of the network’s neurons for when they first fire *≥* 100 Hz. **(C)** Change in the firing rate from after learning to the end of the simulation. **(D)** Weights, *w*, at the end of the simulation. **(E)** Actual potentiation amount as a percentage of that determined by the STDP window. **(F)** Left axis: example of 100 synapses’ weights. Right axis/dashed line: mean of nonfilopodia synapses’ weights. **(G)** Distance between nonfilopodia spines. **(H)** Resource pools, *p*. **(I)** Mean in degree, out degree, in closeness, and out closeness during the simulation. **(J)** Network of neurons after learning. **(K)** Weight change after learning. **(L)** Network of neurons at the end of the simulation. **(J)** and **(L)** Only the strongest 10 % are shown. Line width represents weight.

After learning, weight statistics match those found experimentally—five observations were identified in the introduction:

1. The weight distribution for recurrent connections at the end of the simulation is unimodal, Fig. 2d, matching those observed experimentally.
2. The distribution also shows a large peak of empty synapses: silent synapses.
3. Resource-based STDP endows reduced and nonpotentiability, Fig. 2e. Specifically, the actual potentiation amount as a percentage of that determined by the STDP window function demonstrates that sometimes not enough resources are available to fully potentiate a synapse; some times very little or no potentiation was observed.
4. Resource-based STDP is stable, with weights remaining persistent in a longer simulation, Fig. 3.
5. Weights converge to individual upper bounds despite the positive feedback loop present due to plasticity. It was observed that weights reach their own saturation points, Fig. 2f. Furthermore, due to the distance dependence of heterosynaptic depression, analysis indicates that resource-based heterosynaptic STDP promotes synaptic sparsity and synaptic competition for limited resources, with functional spines becoming spatially distributed along the dendrite. The average distance between a nonfilopodia spine and its nearest nonfilopodia spine neighbor increases during learning, Fig. 2g.

**Figure 3:**
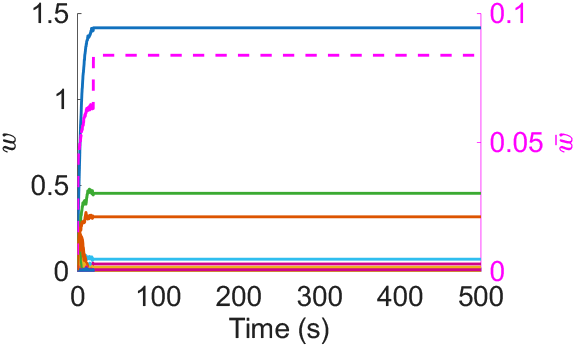
Left axis: example of 100 synapses’ weights during a long simulation. Right axis/dashed line: mean of nonfilopodia synapses’ weights.

Most of the resources available in the neurons’ pools quickly decrease due to plasticity driven changes, Fig. 2h. Some pools may not be used up due to plasticity and instead naturally decay.

During the initial learning phase, the network’s in/out degree increases and stabilizes *≈* 3, and the closeness (where the length of the paths is the axonal delay) decreases, Fig. 2i. The decrease in closeness demonstrates that learning optimizes for short delays.

After the initial learning phase, the network is further optimized with an increase in nonfilopodia spine distance and a decrease in closeness. Furthermore, the network structure changes during this optimization phase with weights increasing or decreasing, Figs. 2j to 2l. Most strong connections are stable with a few changing; the majority of optimization changes are with weaker synapses.

These results in a typical network demonstrate that resource-based STDP is capable of stable learning in recurrent networks with an innate homeostasis mechanism controlling runaway activity and potentiation. Next, to model spine loss in neurodegenerative diseases, synapses are progressively removed, Figs. 4a and 4b.

**Figure 4:**
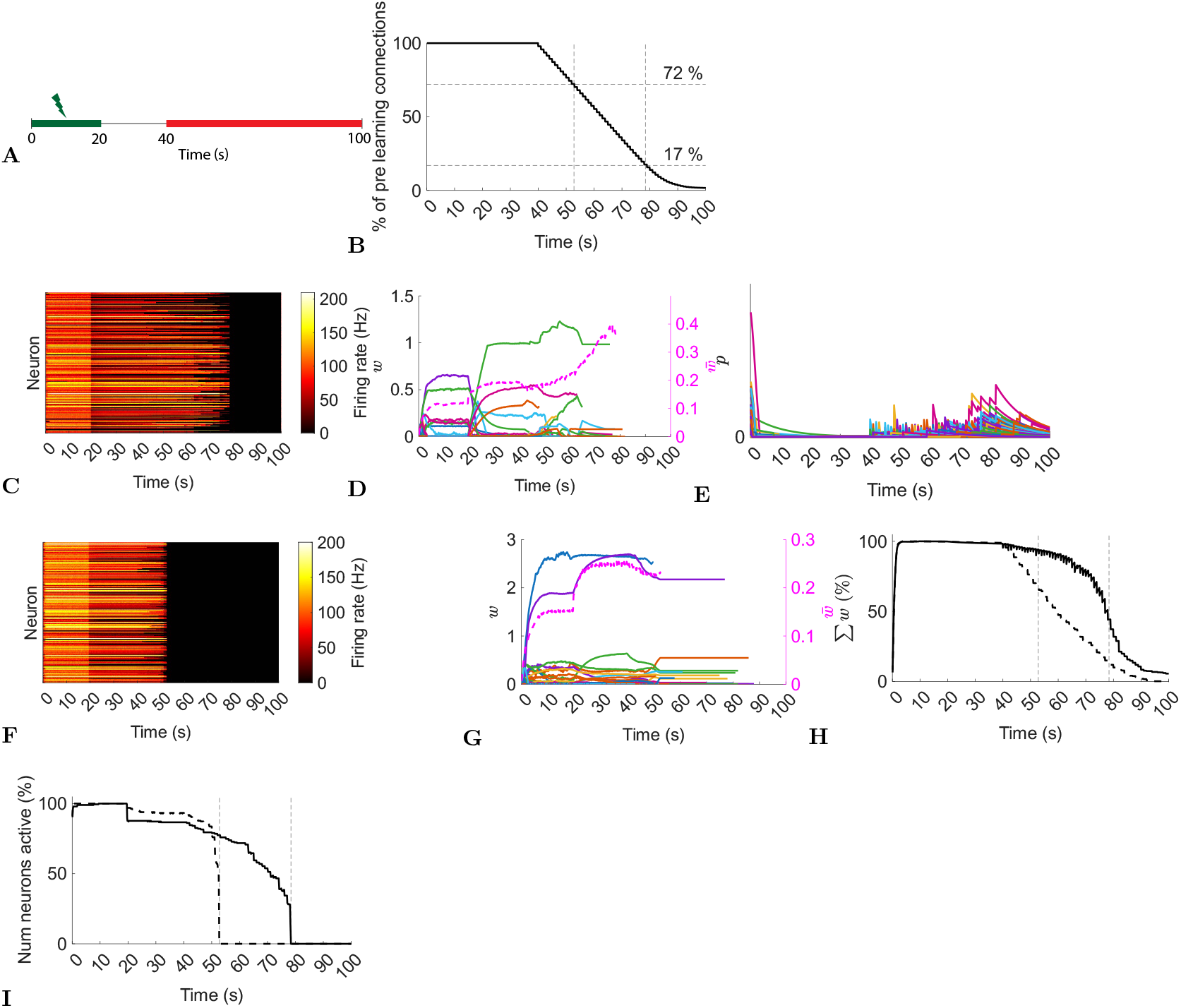
Typical network dynamics with and without resource pool replenishment while removing connections. **(A)** Input is provided for the first 20 s. Synapses are progressively deleted from 40 s on (red). **(B)** Progressive synaptic loss. **(C)** Firing rates of the network’s neurons with replenishment of the resource pool. **(D)** Left axis: example of 100 synapses’ weights with resource pool replenishment. Right axis/dashed line: mean of nonfilopodia synapses’ weights. **(E)** Resource pools, *p*, with replenishment of the resource pool. **(F)** Firing rates of the network’s neurons without replenishment of the resource pool. **(G)** Left axis: example of 100 synapses’ weights without replenishment of the resource pool. Right axis/dashed line: mean of nonfilopodia synapses’ weights. **(H)** Sum of weights as a percentage of the maximum sum of weights. Solid line, with resource replenishment; dashed line, without resource replenishment. **(I)** The number of active neurons as a percentage of the maximum number of active neurons. Solid line: with resource replenishment; Dashed line: without resource replenishment. **(B), (H)**, and **(I)** Vertical lines are when all neurons stopped firing either with or without resource replenishment. **(D)** and **(G)** Weights are shown while they exist and the mean of nonfilopodia synapses’ weights is shown while neurons are active.

In a typical network with resource-based heterosynaptic STDP, the memory is maintained until *≈* 17 % of the original connections remain, Fig. 4c. As synapses are removed, their resources replenish the pool and allow further compensatory potentiation with an increase in mean synaptic weight and transient increases in the pool of resources, Figs. 4d and 4e. However, when this replenishment is blocked, the memory is only maintained until *≈* 72 % synapses remain, Fig. 4f—with no increase in the remaining synaptic weights, Fig. 4g.

This is further illustrated when comparing the sum of weights as a percentage of the maximum sum, Fig. 4h. Under the replenishment-blocking condition, the total synaptic input disappears *≈* 4 times faster than under the control condition. Moreover, loss of neuron activity is sudden without replenishment of resources; whereas with replenishment of resources there is a progressive loss, Fig. 4i.

## 4 Discussion

Discrepancies were reconciled between computational STDP models and empirical observations by using a STDP framework based on resource-dependent heterosynaptic STDP in a recurrent spiking network. By integrating resource-dependent potentiation with heterosynaptic depression at nearby synapses, neurons performed a learning task while preserving synaptic weight attributes and statistical traits consistent with biological data.

Resource-dependent heterosynaptic STDP results in weight distributions characterized by a singular peak and a pronounced tail. The distribution also shows a large peak at zero of spines empty of resources (silent synapses) similar to those estimated by Brunel et al. (2004) and similar to an abundance of filopodia as seen by Yasumatsu et al. (2008) and reviewed by Kasai (2023).

Furthermore, this approach to heterosynaptic STDP incorporates robust competitive dynamics and synaptic homeostasis, leading to varying intrinsic upper limits for individual synapses at which a synapse is no longer potentiable. In addition, the STDP rule promotes sparse spatial encoding in afferents and therefore strong competition between neighboring spines for limited resources. This homeostasis among weights is independent of any additional normalization mechanism or universal constraints to regulate learning. Such synaptic homeostasis is a compelling mechanism for stabilizing neural activity and plasticity within recurrent networks.

Finally, given progressive synaptic loss, resource-based STDP demonstrated an innate compensatory mechanism due to replenishment of resources; this mechanism has been hypothesized to counteract a loss in input (Bhembre et al., 2023). Loss of synaptic input is associated with early stages of neurodegenerative diseases, for example, Alzheimer’s disease (Spires et al., 2005). Resource-based STDP demonstrated the ability to maintain total synaptic input by replenishing resources in the pool after synaptic removal. Moreover, resource-based STDP demonstrated a sufficient substrate for observations showing enlargement of synapses following insults such as deafferentation and sensory deprivation (Chen and Hillman, 1982; Barnes et al., 2017) and the hypothesized spine enlargement in Alzheimer’s disease (Bhembre et al., 2023). Finally, resource-based STDP exhibited a progressive loss in, rather than a sudden loss in, neuron activity. How this relates to aberrant activity observed in Alzheimer’s disease (Korzhova et al., 2021) is unknown, and exploring this offers an interesting future extension to this work.

Given that the proposed STDP rule aligns with experimentally observed weight statistics and supports a hypothesized neurodegenerative compensatory mechanism, its potential biological plausibility warrants examination. Royer et al. (2003) observed in amygdala slices that the induction of long-term depression (potentiation) resulted in corresponding long-term potentiation (depression) at distally located dendritic sites, determined by the distance from the initial induction site. In contrast, Hou et al. (2008) reported no such synaptic changes at adjacent sites in cultured hippocampal neurons derived from rat embryos.

The research carried out by Oh et al. (2015) in P6-P7 hippocampal slice cultures aligns with the proposed STDP model. Stimulation-induced potentiation in specific spines was demonstrated to cause a reduction in the size of adjacent unstimulated spines. This observation can be explained by two potential mechanisms: first, the competition among nearby spines for a scarce resource, or second, an activity-triggered signal facilitating the reduction of adjacent spines. The findings of Oh et al. lend credence to the second hypothesis. They determined that blocking calcineurin, IP_3_ receptors or group I metabotropic glutamate receptors hinders heterosynaptic shrinkage, without affecting the potentiation process when Ca^2+^/calmodulin-dependent protein kinase II (CaMKII) is inhibited. This evidence indicates that synaptic potentiation and the associated decrease in nearby synaptic strength operate through distinct pathways.

Bian et al. (2015) observed that competition among spines for cadherin/catenin complexes plays a crucial role in orchestrating both the maturation of individual spines and the pruning of adjacent spines. Their *in vivo* studies revealed that variations in cadherin/catenin complex concentrations between neighboring spines lead to a redistribution of *β*-catenin, which in turn influences whether a spine matures or is trimmed. Crucially, they demonstrated that this process depends on neuronal activity and the distance between the enlarging spine and the nearby one that is slated to be eliminated. Furthermore, this competitive mechanism was not limited to individual neurons; it occurred in neighboring neurons receiving similar axonal input.

Within the framework of long-term potentiation (LTP), the relocation of AMPA receptors to synaptic sites is associated with increased synaptic activity (Hayashi et al., 2000; Sutton and Schuman, 2006; Shi et al., 2001). In this process, CaMKII experiences autophosphorylation when intracellular Ca^2+^ levels rise through NMDA receptor-mediated channels, culminating in the phosphorylation of GluR1 (Roche et al., 1996). AMPA receptors can originate from multiple sources, such as recycling endosomes (Park et al., 2004) and the trans-Golgi network (Horton and Ehlers, 2004). Our model consolidates these different origins into a common pool, taking advantage of evidence that AMPA receptors can traverse long distances, facilitated by movement along dendritic membranes (Choquet and Triller, 2003) and microtubule pathways (Washbourne et al., 2002). Regarding the mechanism that underlies heterosynaptic depression, Oh et al. (2015) observed that blocking calcineurin, IP_3_Rs, or group I metabotropic glutamate receptors hindered the contraction of adjacent spines. Additionally, Bian et al. (2015) propose the existence of a single molecular structure that fulfills dual roles, both as a ‘resource’ and as a transmitter of depressive signals. Finally, recent experimental and computational evidence suggests that Ca^2+^ activity is a key component of competitive heterosynaptic plasticity (Chater et al., 2024).

The depiction of STDP herein does not include explicit modeling of biological signaling pathways or diffusible molecules that might cause depression in neighboring synapses after activity-driven potentiation, nor does it specify a precise timeline for such processes. Furthermore, it does not incorporate the transfer of resources into or out of a dendritic spine. Introducing these complexities could impose interesting limitations on the model, suggesting areas for future investigation.

Although a precise ‘resource’ is not identified, nor is a specific depression signal characterized, multiple hypotheses suitable for experimental examination can be proposed under healthy and synaptic degenerative conditions.

- As described by Oh and colleagues (2015), suppressing the heterosynaptic depression signal could allow learning driven by potentiation under healthy conditions to continue until resources in the pool(s) are exhausted. Subsequent learning would require the creation of new resources. As a result, while the pace of learning may reduce under signal-suppressed conditions compared to when active signals are present, it would not come to a complete stop until all resources are used with no change in neighboring spines.
- Suppression of the depression signal under synaptic degenerative conditions might interfere with memory retention during synaptic loss if the signal is required for resource replenishment.
- Excessive activation of the depression signal in both healthy and synaptic degenerative conditions could result in a temporary surplus of resources that would persist until they are used by potentiating synapses or degrade over time.
- A reduction in total resources could hinder learning under healthy conditions, although it would not stop if vital resources are actively liberated through heterosynaptic depression.
- A reduction in total resources under synaptic degenerative conditions could accelerate the total loss of synaptic input and therefore the loss of neuronal/memory activity.
- In contrast, an increase in available resources might lead to uncontrolled neuronal activity and potentiation under health conditions.
- Under conditions of synaptic degeneration, an increase in available resources can contribute to memory retention.

## References

Abraham, W. C., Logan, B., Greenwood, J. M., and Dragunow, M. (2002). Induction and Experience-Dependent Consolidation of Stable Long-Term Potentiation Lasting Months in the Hippocampus. The Journal of Neuroscience 22, 9626–9634. doi:10.1523/jneurosci.22-21-09626.2002

Bannon, N. M., Chistiakova, M., and Volgushev, M. (2020). Synaptic Plasticity in Cortical Inhibitory Neurons: What Mechanisms May Help to Balance Synaptic Weight Changes? Frontiers in Cellular Neuroscience 14, 204. doi:10.3389/fncel.2020.00204

Barbour, B., Brunel, N., Hakim, V., and Nadal, J.-P. (2007). What can we learn from synaptic weight distributions? Trends in Neurosciences 30, 622–629. doi:10.1016/j.tins.2007.09.005

Barnes, S. J., Franzoni, E., Jacobsen, R. I., Erdelyi, F., Szabo, G., Clopath, C., et al. (2017). Deprivation-Induced Homeostatic Spine Scaling InVivo Is Localized to Dendritic Branches that Have Undergone Recent Spine Loss. Neuron 96, 871–882.e5. doi:10.1016/j.neuron.2017.09.052

Bhembre, N., Bonthron, C., and Opazo, P. (2023). Synaptic Compensatory Plasticity in Alzheimers Disease. The Journal of Neuroscience 43, 6833–6840. doi:10.1523/jneurosci.0379-23.2023

Bi, G.-q. and Poo, M.-m. (1998). Synaptic Modifications in Cultured Hippocampal Neurons: Dependence on Spike Timing, Synaptic Strength, and Postsynaptic Cell Type. The Journal of Neuroscience 18, 10464–10472. doi:10.1523/jneurosci.18-24-10464.1998

Bian, W.-J., Miao, W.-Y., He, S.-J., Qiu, Z., and Yu, X. (2015). Coordinated Spine Pruning and Maturation Mediated by Inter-Spine Competition for Cadherin/Catenin Complexes. Cell 162, 808–822. doi:10.1016/j.cell.2015.07.018

Billings, G. and Rossum, M. C. W. v. (2009). Memory Retention and Spike-Timing-Dependent Plasticity. Journal of Neurophysiology 101, 2775–2788. doi:10.1152/jn.91007.2008

Brunel, N., Hakim, V., Isope, P., Nadal, J.-P., and Barbour, B. (2004). Optimal Information Storage and the Distribution of Synaptic Weights Perceptron versus Purkinje Cell. Neuron 43, 745–757. doi:10.1016/j.neuron.2004.08.023

Buzski, G. (2004). Large-scale recording of neuronal ensembles. Nature Neuroscience 7, 446–451. doi:10.1038/nn1233

Buzski, G. and Mizuseki, K. (2014). The log-dynamic brain: how skewed distributions affect network operations. Nature Reviews Neuroscience 15, 264–278. doi:10.1038/nrn3687

Chater, T. E., Eggl, M. F., Goda, Y., and Tchumatchenko, T. (2024). Competitive processes shape multi-synapse plasticity along dendritic segments. Nature Communications 15, 7572. doi:10.1038/s41467-024-51919-0

Chater, T. E. and Goda, Y. (2021). My Neighbour Heterodeconstructing the mechanisms underlying heterosynaptic plasticity. Current Opinion in Neurobiology 67, 106–114. doi:10.1016/j.conb.2020.10.007

Chen, J. Y., Lonjers, P., Lee, C., Chistiakova, M., Volgushev, M., and Bazhenov, M. (2013). Heterosynaptic Plasticity Prevents Runaway Synaptic Dynamics. Journal of Neuroscience 33, 15915–15929. doi:10.1523/jneurosci.5088-12.2013

Chen, S. and Hillman, D. E. (1982). Plasticity of the parallel Fiber-Purkinje cell synapse by spine takeover and new synapse formation in the adult rat. Brain Research 240, 205–220. doi:10.1016/0006-8993(82)90217-7

Choquet, D. and Triller, A. (2003). The role of receptor diffusion in the organization of the postsynaptic membrane. Nature Reviews Neuroscience 4, 251–265. doi:10.1038/nrn1077

Cooper, L. N. and Bear, M. F. (2012). The BCM theory of synapse modification at 30: interaction of theory with experiment. Nature Reviews Neuroscience 13, 798–810. doi:10.1038/nrn3353

Debanne, D., Ghwiler, B. H., and Thompson, S. M. (1999). Heterogeneity of Synaptic Plasticity at Unitary CA3CA1 and CA3CA3 Connections in Rat Hippocampal Slice Cultures. The Journal of Neuroscience 19, 10664–10671. doi:10.1523/jneurosci.19-24-10664.1999

Dong, Z., Han, H., Li, H., Bai, Y., Wang, W., Tu, M., et al. (2015). Long-term potentiation decay and memory loss are mediated by AMPAR endocytosis. Journal of Clinical Investigation 125, 234–247. doi:10.1172/jci77888

Eckmann, S., Young, E. J., and Gjorgjieva, J. (2024). Synapse-type-specific competitive Hebbian learning forms functional recurrent networks. Proceedings of the National Academy of Sciences 121, e2305326121. doi:10.1073/pnas.2305326121

Hayashi, Y., Shi, S.-H., Esteban, J. A., Piccini, A., Poncer, J.-C., and Malinow, R. (2000). Driving AMPA Receptors into Synapses by LTP and CaMKII: Requirement for GluR1 and PDZ Domain Interaction. Science 287, 2262–2267. doi:10.1126/science.287.5461.2262

Herms, J. and Dorostkar, M. M. (2015). Dendritic Spine Pathology in Neurodegenerative Diseases. Annual Review of Pathology: Mechanisms of Disease 11, 1–30. doi:10.1146/annurev-pathol-012615-044216

Horton, A. C. and Ehlers, M. D. (2004). Secretory trafficking in neuronal dendrites. Nature Cell Biology 6, 585–591. doi:10.1038/ncb0704-585

Hou, Q., Zhang, D., Jarzylo, L., Huganir, R. L., and Man, H.-Y. (2008). Homeostatic regulation of AMPA receptor expression at single hippocampal synapses. Proceedings of the National Academy of Sciences 105, 775–780. doi:10.1073/pnas.0706447105

Humble, J. (2013). Learning, self-organisation and homeostasis in spiking neuron networks using spike-timing dependent plasticity. Ph.D. thesis

Humble, J., Hiratsuka, K., Kasai, H., and Toyoizumi, T. (2019). Intrinsic Spine Dynamics Are Critical for Recurrent Network Learning in Models With and Without Autism Spectrum Disorder. Frontiers in Computational Neuroscience 13, 38. doi:10.3389/fncom.2019.00038

Hunt, S., Leibner, Y., Mertens, E. J., Barros-Zulaica, N., Kanari, L., Heistek, T. S., et al. (2022). Strong and reliable synaptic communication between pyramidal neurons in adult human cerebral cortex. Cerebral Cortex 33, 2857–2878. doi:10.1093/cercor/bhac246

Kasai, H. (2023). Unraveling the mysteries of dendritic spine dynamics: Five key principles shaping memory and cognition. Proceedings of the Japan Academy. Series B, Physical and Biological Sciences 99, 254–305. doi: 10.2183/pjab.99.018

Kim, H. J., Lee, S., Kim, G. H., Sung, K., Yoo, T., Pyo, J. H., et al. (2025). GluN2B-mediated regulation of silent synapses for receptor specification and addiction memory. Experimental & Molecular Medicine, 1–14 doi: 10.1038/s12276-025-01399-z

Korzhova, V., Marinkovi, P., Njavro, J. R., Goltstein, P. M., Sun, F., Tahirovic, S., et al. (2021). Long-term dynamics of aberrant neuronal activity in awake Alzheimers disease transgenic mice. Communications Biology 4, 1368. doi:10.1038/s42003-021-02884-7

Lemarchal, J.-D., Jedynak, M., Trebaul, L., Boyer, A., Tadel, F., Bhattacharjee, M., et al. (2021). A brain atlas of axonal and synaptic delays based on modelling of cortico-cortical evoked potentials. Brain 145, 1653–1667. doi:10.1093/brain/awab362

Meftah, S. and Gan, J. (2023). Alzheimers disease as a synaptopathy: Evidence for dysfunction of synapses during disease progression. Frontiers in Synaptic Neuroscience 15, 1129036. doi:10.3389/fnsyn.2023.1129036

Montgomery, J. M. and Madison, D. V. (2004). Discrete synaptic states define a major mechanism of synapse plasticity. Trends in Neurosciences 27, 744–750. doi:10.1016/j.tins.2004.10.006

Morrison, A., Aertsen, A., and Diesmann, M. (2007). Spike-Timing-Dependent Plasticity in Balanced Random Networks. Neural Computation 19, 1437–1467. doi:10.1162/neco.2007.19.6.1437

Morrison, A., Diesmann, M., and Gerstner, W. (2008). Phenomenological models of synaptic plasticity based on spike timing. Biological Cybernetics 98, 459–478. doi:10.1007/s00422-008-0233-1

Oh, W., Parajuli, L., and Zito, K. (2015). Heterosynaptic Structural Plasticity on Local Dendritic Segments of Hippocampal CA1 Neurons. Cell Reports 10, 162–169. doi:10.1016/j.celrep.2014.12.016

Park, M., Penick, E. C., Edwards, J. G., Kauer, J. A., and Ehlers, M. D. (2004). Recycling Endosomes Supply AMPA Receptors for LTP. Science 305, 1972–1975. doi:10.1126/science.1102026

Penzes, P., Cahill, M. E., Jones, K. A., VanLeeuwen, J.-E., and Woolfrey, K. M. (2011). Dendritic spine pathology in neuropsychiatric disorders. Nature Neuroscience 14, 285–293. doi:10.1038/nn.2741

Rall, W. (1969). Time Constants and Electrotonic Length of Membrane Cylinders and Neurons. Biophysical Journal 9, 1483–1508. doi:10.1016/s0006-3495(69)86467-2

Roche, K. W., O’Brien, R. J., Mammen, A. L., Bernhardt, J., and Huganir, R. L. (1996). Characterization of Multiple Phosphorylation Sites on the AMPA Receptor GluR1 Subunit. Neuron 16, 1179–1188. doi:10.1016/s0896-6273(00)80144-0

Rossum, M. C. W. v., Bi, G. Q., and Turrigiano, G. G. (2000). Stable Hebbian Learning from Spike Timing-Dependent Plasticity. The Journal of Neuroscience 20, 8812–8821. doi:10.1523/jneurosci.20-23-08812.2000

Royer, S. and Par, D. (2003). Conservation of total synaptic weight through balanced synaptic depression and potentiation. Nature 422, 518–522. doi:10.1038/nature01530

Shi, S.-H., Hayashi, Y., Esteban, J. A., and Malinow, R. (2001). Subunit-Specific Rules Governing AMPA Receptor Trafficking to Synapses in Hippocampal Pyramidal Neurons. Cell 105, 331–343. doi:10.1016/s0092-8674(01)00321-x

Spires, T. L., Meyer-Luehmann, M., Stern, E. A., McLean, P. J., Skoch, J., Nguyen, P. T., et al. (2005). Dendritic Spine Abnormalities in Amyloid Precursor Protein Transgenic Mice Demonstrated by Gene Transfer and Intravital Multiphoton Microscopy. The Journal of Neuroscience 25, 7278–7287. doi:10.1523/jneurosci.1879-05.2005

Sutton, M. A. and Schuman, E. M. (2006). Dendritic Protein Synthesis, Synaptic Plasticity, and Memory. Cell 127, 49–58. doi:10.1016/j.cell.2006.09.014

Turrigiano, G. G. (2008). The Self-Tuning Neuron: Synaptic Scaling of Excitatory Synapses. Cell 135, 422–435. doi:10.1016/j.cell.2008.10.008

Washbourne, P., Bennett, J. E., and McAllister, A. K. (2002). Rapid recruitment of NMDA receptor transport packets to nascent synapses. Nature Neuroscience 5, 751–759. doi:10.1038/nn883

Yasumatsu, N., Matsuzaki, M., Miyazaki, T., Noguchi, J., and Kasai, H. (2008). Principles of Long-Term Dynamics of Dendritic Spines. The Journal of Neuroscience 28, 13592–13608. doi:10.1523/jneurosci.0603-08.2008

