## Supplementary materials for "Resource-dependent heterosynaptic spike-timing-dependent plasticity in recurrent networks with and without synaptic degeneration"

### Supplementary Material

#### 1 METHODS

##### 1.1 Additive STDP

Equation S1 describes the STDP function used. The learning rate $A = 0.2$ was assigned so that the learning was slow. Recurrent weights were bounded $[0, 1]$.

$$f(\tau) = \begin{cases} A \exp\left(\frac{-\tau}{\tau_{STDP}}\right) & \text{if } \tau \geq 0 \\ \\ A \exp\left(\frac{\tau}{\tau_{STDP}}\right) & \text{if } \tau < 0 \end{cases} \tag{S1}$$

##### 1.2 Multiplicative STDP

Equations S2 and S3 describe the STDP function used where $w$ is the weight that is being updated. The learning rate $A = 0.05$ was assigned so that the learning was slow. Recurrent weights were bounded $[0, 1]$.

$$f(\tau) = \begin{cases} A \exp\left(\frac{-\tau}{\tau_{STDP}}\right) & \text{if } \tau \geq 0 \\ \\ A \exp\left(\frac{\tau}{\tau_{STDP}}\right) \; w & \text{if } \tau < 0 \end{cases} \tag{S2}$$

$$f(\tau) = \begin{cases} A \exp\left(\frac{-\tau}{\tau_{STDP}}\right) & \text{if } \tau \geq 0 \\ \\ A \exp\left(\frac{\tau}{\tau_{STDP}}\right) \; \frac{w}{0.5} & \text{if } \tau < 0 \end{cases} \tag{S3}$$

#### 2 RESULTS

When using additive STDP or multiplicative STDP (Eqs. S1 and S2/S3 respectively) activity increases and the weights converge to unimodal distributions, Fig. 1. In the additive case, potentiation leads to increased activity, and increased activity leads to potentiation; this results in a unimodal distribution at the upper bound. In the multiplicative case, the distribution mean depends on the weight dependence: if weak, it is similar to additive, and if stronger, the weights converge to the weight dependence value. Under additive and multiplicative STDP, all neurons are recruited, and all fire at the maximum firing rate allowed under the refractory constraint.

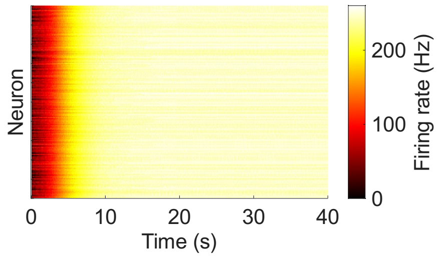


**Figure 1a.**

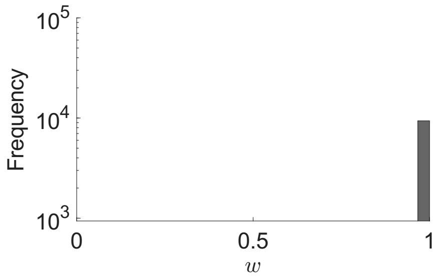


**Figure 1b.**

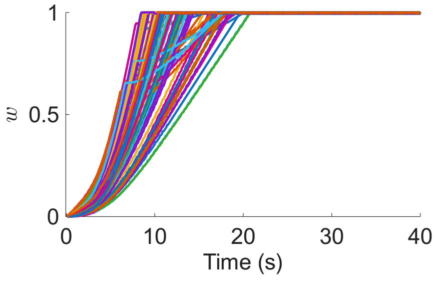


**Figure 1c.**

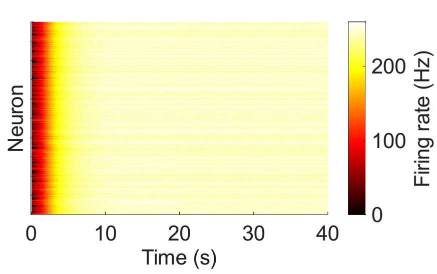


**Figure 1d.**

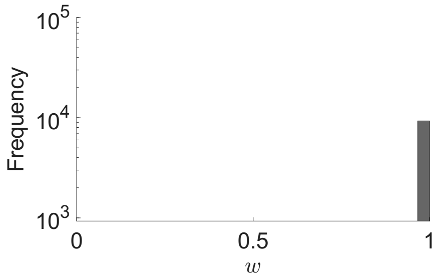


**Figure 1e.**

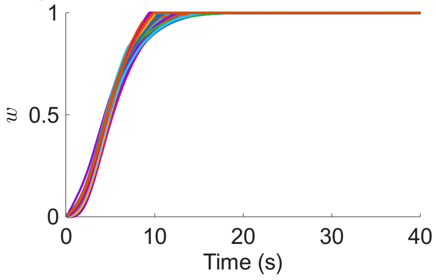


**Figure 1f.**

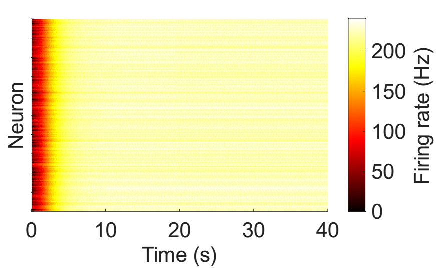


**Figure 1g.**

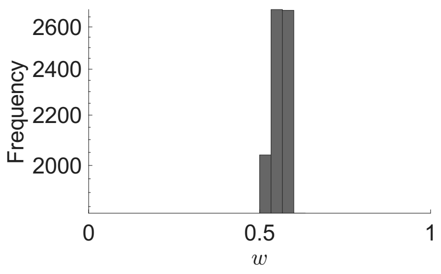


**Figure 1h.**

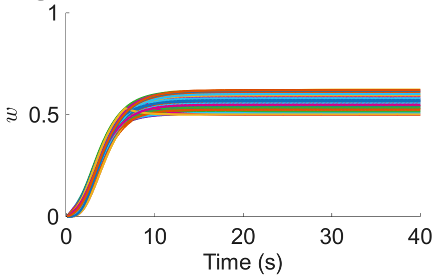


**Figure 1i.**

**Figure 1.** Typical network dynamics with additive or multiplicative STDP. **(A)** Firing rates of the network's neurons with additive STDP. **(B)** Weights, $w$, after learning with additive STDP. **(C)** Example of $100$ synapses' weights with additive STDP. **(D-F)** Same as A-C, but with multiplicative STDP with Eq. S2 **(G-I)** Same as A-C, but with multiplicative STDP with Eq. S3
